# Bimodal breeding phenology in the Parsley Frog *Pelodytes punctatus* as a bet-hedging strategy in an unpredictable environment despite strong priority effects

**DOI:** 10.1101/2022.02.24.481784

**Authors:** Hélène Jourdan-Pineau, Pierre-André Crochet, Patrice David

**Affiliations:** CIRAD, UMR ASTRE, F-34398 Montpellier, France. ASTRE, Univ Montpellier, CIRAD, INRAE, Montpellier, France; CEFE, UMR 5175 CNRS, Montpellier, France

**Keywords:** breeding phenology, bet-hedging, priority effects, stochastic environment, anuran

## Abstract

When environmental conditions are unpredictable, expressing alternative phenotypes spreads the risk of failure, a mixed strategy called bet-hedging. In the southern part of its range, the Parsley Frog *Pelodytes punctatus* breeds both in autumn and in spring. Our aim was to study the breeding phenology and reproductive success associated with the use of those two seasonal niches to understand how this breeding strategy can be maintained. Field surveys revealed that breeding phenology was typically bimodal with a higher breeding effort in autumn. More importantly, in spring, the survival rate of offspring was severely reduced by the presence of autumn tadpoles, indicating a clear priority effect. However, the autumn cohort often failed to survive over winter, in which case spring cohorts were often successful. Based on those results, we constructed a model in which females can allocate a variable portion of eggs to each season and added a priority effect. We conclude that the existence of the two breeding seasons may indeed constitute a bet-hedging strategy.

## Introduction

Breeding phenology is one of the key components of adaptation to temporally variable environments. Temporal dynamics of both the biotic and abiotic environment impose selective constraints on parental development and physiological state (to be able to reproduce) as well as offspring survival (at the various developmental stages until they reach sexual maturity and start to reproduce) (Rand, 1973). There is a vast amount of literature on intraspecific variation of breeding patterns in relation to environmental conditions, in particular latitude, altitude and climate. In the context of current climate change, many species in temperate regions have advanced their breeding time (e.g. Brown et al., 2016; Frederiksen et al., 2004; Møller, 2008), as a result of microevolutionary changes and/or of phenotypic plasticity (Charmantier & Gienapp, 2014). Most of these studies concern species with a single reproductive peak in the year, which has to match as precisely as possible a seasonal peak of resource availability in order to maximize reproductive success (e.g. caterpillar availability for tits). The exact date of the resource peak may vary from year to year and species usually rely on cues to anticipate it and plastically delay or advance the onset of reproduction every year. However, in some cases reproductive success depends on even more irregular and/or unpredictable conditions. In such situations, species face the risk of complete reproductive failure at any given breeding attempt, a regime that favors the expression of alternative phenotypes to spread the risk (Cohen, 1970; Slatkin, 1974; Philippi & Seger, 1989; Leimar, 2005; Venable, 2007).

Theory predicts that in stochastic environments, selection favors life history traits that reduce temporal fitness variation even if they result in lowered arithmetic mean fitness (Philippi & Seger, 1989). This risk-spreading strategy is called bet-hedging. In temporally variable environments, the long-term fitness of a genotype is measured by the geometric mean of the fitness contribution over successive generations for a particular genotype (Lewontin & Cohen, 1969; Olofsson et al., 2009; Simons, 2011; Yasui & Yoshimura, 2018). This geometric mean fitness is highly impacted by low values; hence, traits with lower fitness variation may have higher long-term fitness. In principle, lower variation in fitness can be achieved either using the same low-risk strategy (conservative bet-hedging), or displaying several strategies, either at once or over several instances (diversified bet-hedging). While there is an abundant theoretical literature on bet-hedging, empirical studies have provided limited evidence so far (Simons, 2011), and the most comprehensive examples concern the timing of germination/diapause and the fraction of dormant seeds/diapausing eggs (Venable, 2007; Gremer et al., 2016; García-Roger et al., 2017; Wang & Rogers, 2018). Some studies even report experimental evolution of bet-hedging traits in response to unpredictable environment (Beaumont et al., 2009; Graham et al., 2014; Maxwell & Magwene, 2017; Tarazona et al., 2017).

One of the best examples of stochastic, unpredictable environments is provided by temporary ponds, alternating between inundation and drought where each breeding event is a bet as habitat desiccation can occur before the end of the breeding cycle. Several examples of bet-hedging occur in temporary ponds. Fairy shrimps (Anostraca) produce drought-resistant eggs showing asynchronous hatching at different hydroperiods (Saiah & Perrin, 1990; Simovich & Hathaway, 1997; Wang & Rogers, 2018). Similarly, rotifers produce diapausing eggs to overpass unfavorable planktonic growing season and only a fraction of those eggs hatch when conditions are suitable (García-Roger et al., 2017; Tarazona et al., 2017).

For amphibian species breeding in temporary ponds, drought can cause 100% mortality of eggs or larvae, resulting in complete failure of one breeding event. One way to reduce the risk of losing a breeding opportunity entirely is to spread this risk at a spatial scale, partitioning broods into various pools as done by the Neotropical poison frog *Allobates femoralis* (Erich et al., 2015). Another bet-hedging strategy could be to split the breeding effort at a temporal scale and exploit all suitable temporal windows.

From an ecological point of view, such temporal niche partitioning is expected to reduce inter and intra-specific competition as well as resource depletion (Carothers et al., 1984). For example, species may share the same habitat but have opposite activity patterns (nocturnal versus diurnal species), as is the case in Neotropical felid community or in grassland ants (Albrecht & Gotelli, 2001; Di Bitetti et al., 2010). The same type of temporal segregation at a daily scale is also observed within species: brown trout *Salmo trutta* reduces competition for a limiting resource by sequential use of foraging areas (Alanärä et al., 2001). Voltinism in insects is another well-studied example of temporal partitioning at the annual scale which is an adaptation to predictable seasonal cycles (Kivelä et al., 2013; Zeuss et al., 2017; Forrest et al., 2019).

However, if successive seasonal cohorts overlap, fitness gains may be asymmetric, because progeny produced by late breeding may suffer from competition or even predation from earlier cohorts (Morin, 1987; Ryan & Plague, 2004; Eitam et al., 2005). Those priority effects are often difficult to disentangle from seasonal effects due to environmental differences experienced by the temporal cohorts (Morin et al., 1990). If priority effects are strong, late breeders may select breeding sites in order to limit the competition by conspecifics (Halloy & Fiaño, 2000; Halloy, 2006; Sadeh et al., 2009) and this may restrict late breeders to poorer sites (Crump, 1991).

In amphibians, this temporal partitioning of breeding activity is thought to regulate community dynamics through interspecific competition (Lawler & Morin, 1993; Gottsberger & Gruber, 2004; Richter-Boix et al., 2006a, 2007c). Similarly, community composition may depend on species arrival and priority effects whereby species arriving earlier monopolize available resources and gain a competitive advantage over late species (Morin et al., 1990; Blaustein & Margalit, 1996; Urban & De Meester, 2009). In Mediterranean regions, climatic conditions are characterized by dry, hot summers and mild winters, with the maximum rainfalls in autumn and spring. This leads to large breeding asynchrony observed between and within amphibian species (Diaz-Paniagua 1988; Jakob, Poizat et al. 2003; Richter-Boix, Llorente et al. 2006; Vignoli, Bologna et al. 2007): whereas most species typically breed in spring, some species breed earlier at the end of winter, and some even breed in autumn in addition to spring.

The Parsley Frog *Pelodytes punctatus* is a small sized anuran distributed in Spain and most of France (locally reaching neighboring countries). It has a broad ecological niche but has poor competitive abilities and is sensitive to fish predation (Morand & Pierre, 1995; Crochet et al., 2004; Richter-Boix et al., 2007b); it thus prefers seasonally flooded habitats to large permanent water bodies (Guyétant et al., 1999; Salvidio et al., 2004; Richter-Boix et al., 2007a). In Spain the Parsley Frog shows a bimodal breeding pattern with higher reproductive effort in spring than in autumn (Guyétant et al., 1999; Richter-Boix et al., 2006a). In France, in addition to spring breeding, autumnal breeding is also observed in Mediterranean regions and areas with mild oceanic climate (Guyétant et al., 1999; Jakob et al., 2003; Richter-Boix et al., 2006b; Cayuela et al., 2012) but the importance of autumn versus spring reproduction has not been quantified. In the rest of the range and at higher altitudes, only spring breeding occurs.

In the Mediterranean areas of southern France, the Parsley Frog uses temporary ponds that refill in September and October but may dry during autumn or later in late spring. Adults thus have to deal with very unpredictable environmental conditions for their future offspring. In addition to this unpredictable risk, tadpoles hatched in autumn or spring are exposed to very different environmental conditions, the most obvious being that the autumn tadpoles overwinter while the spring ones do not. This should result in drastically different developmental trajectories but also in different offspring survivals. Both seasonal cohorts may also interact, leading to a potential competitive advantage to the earlier cohort over the later, i.e. a priority effect. The relative success of each breeding period and the outcome of the interaction between cohorts are key parameters to understand the persistence of this two-peaks breeding strategy. In fact, several pieces of information are still lacking in order to understand the evolutionary basis of this seasonally variable breeding strategy. Do we have a single protracted breeding season or a really bimodal reproduction generated by the coexistence of alternative breeding timing? If so, what is the relative importance of autumn versus spring reproduction? What is the survival of offspring produced at the two breeding periods and how is it affected by the presence of conspecifics? Once this basic knowledge is obtained, it can be fed into theoretical models for the evolution of mixed breeding strategies.

In this paper, we characterize the breeding phenology (temporal dynamic, relative proportion of each breeding period) of Parsley Frog in a French Mediterranean area based on results from a 3-year field survey. We monitored the survival of offspring produced in each season to estimate the success of this breeding strategy. We also investigated the factors influencing breeding and tadpole survival, in particular whether there is a priority effect between seasonal cohorts. Finally, using an analytical model adapted from Cohen (Cohen, 1966) we tested whether the coexistence of the two breeding periods can be interpreted as a bet-hedging strategy.

## Methods

### Field survey

The field study was carried out from September 2007 to August 2010 in 19 ponds situated around Montpellier, southern France (Annex 1). Those ponds are man-made environments, often dug out to provide drinking water for livestock (sheep and cows) or for game. The ponds surveyed included temporary and permanent sites. We define here the autumn breeding season as the period spanning from September to December and the spring breeding season from January to April. We surveyed each pond twice each month. During each visit, we recorded the depth of the pond.

### Sampling methods

At every visit (mostly diurnal), we looked for newly deposited egg masses throughout the entire water body and classified the egg masses as small, medium and large, corresponding to an average of 75, 150 and 250 eggs per mass, respectively (Salvador & Paris 2001, and personal observation). The Parsley Frog’s embryonic period ranges from 5 days at 15°C to 15 days at 10°C (Toxopeus et al., 1993). Moreover, embryos stay attached to the jelly for several additional days (Guyétant et al., 1999). Thus, with an interval of 15 days between two successive visits, we may have missed a few masses but we have avoided double-counting masses since 15-day old masses can readily be distinguished from new ones based on the developmental stages of the embryos. In only 2% of the larval cohorts produced were small larvae observed in ponds where we did not notice the presence of egg masses before. The probability of detection of an egg mass, even if not perfect, was similar in autumn and in spring.

We estimated the number of amphibian larvae and invertebrates present in the ponds using 5 to 10 dipnet sweeps (depending on the pond size). The anuran community of the area consists of 7 species: *Pelodytes punctatus, Pelobates cultripes, Alytes obstetricans, Bufo bufo, Epidalea calamita, Hyla meridionalis*, and *Pelophylax sp., (P. ridibundus* and/or *P. perezi* & *P*. kl. *grafi*, depending on the sites). Potential predators of tadpoles are urodeles and aquatic invertebrates. Two urodele species (*Lissotriton helveticus* and *Triturus marmoratus*) were recorded in the ponds but due to the rare occurrence of *Triturus marmoratus*, only *Lissotriton helveticus* was included in subsequent analyses (as adults as well as larvae).

We also surveyed dragonfly larvae (Anisoptera) and backswimmers (Heteroptera, Notonectidae) that are potential predators of tadpoles (Richter-Boix et al., 2007a) except during the first year. Diving beetles (Coleoptera, Dytiscidae) are also known to prey on tadpoles but were very rare in the studied ponds and thus not considered for this study.

We divided the total counts for each amphibian larvae and invertebrate predators captured in each pond by the number of dipnet sweeps taken in each pond. This procedure yielded a crude proxy for density on the basis of catch per unit effort and could therefore be compared across localities.

### Reproductive effort and offspring survival

Reproductive strategy of Parsley Frog was described by two measures: the *presence of egg masses* (binary variable: whether some eggs were laid or not when we visited a pond) and the *number of egg masses* (integer, non-zero; applies only to cases where egg masses are present). We normalized the number of egg masses by their size (e.g. a small egg mass equals ½ medium egg mass).

For each breeding event, we estimated the *hatching rate* as the ratio of the number of small tadpoles (Gosner stage 26, free swimming tadpole) to the number of eggs spawned. Similarly, we quantified the *survival rate from egg to metamorph* as the ratio of the number of metamorphs (Gosner stage 42-43) to the number of eggs spawned. Finally, we calculated the *survival rate during larval stage* as the ratio of the number of metamorphs over the number of small tadpoles. This index could only be estimated in about one third of the breeding events when hatching was successful (i.e. the number of small tadpoles was not null).

The number of tadpoles in a pond was estimated using the mean number of tadpoles caught per dipnet sweep scaled to a sampling surface of 1 m^2^ (we estimated that one dipnet sweep sampled a surface of 0.5 m^2^, taking the dipnet size and the length of the haul into account) and then multiplied by the surface of the pond. This should not be taken as an attempt to estimate precisely the number of tadpoles present in a pond at a given time but as an index of abundance that can be compared between ponds and between breeding events. It was sometimes impossible to follow the larval development and metamorphosis of offspring from a particular breeding event. Indeed, Parsley Frogs may breed three to four times during each seasonal breeding event. In these cases, the successive sub-cohorts produced are indistinguishable after a few weeks, and we summed the eggs counted in two or three successive visits to evaluate survival from a combination of breeding events within a given season (and within a site). Survival measures should be viewed as an index to assess the differences of reproductive success between seasons as there is no reason to expect any seasonal bias in this index.

### Explanatory variables

Explanatory variables for the breeding probability and breeding effort are the season, depth of the pond as well as the presence of conspecific and inter-specific competitors (larvae of anuran species) and predators (invertebrates and adult newts) in the pond. Except for the depth of the pond, all those explanatory variables were also applied to explain the success (offspring survival) of breeding events. We summed the density of competitors and similarly the density of predators despite the differences in competitive performance and predation pressure of the various species toward Parsley Frog tadpoles.

To assess the potential impact of predation and competition on survival rates, we evaluated the mean density of predators and competitors encountered by Parsley Frog tadpoles during their larval development. More precisely, data from literature indicates that only small tadpoles (<12 mm snout-vent length) have lower survival due to predation by aquatic invertebrates (Tejedo, 1993). Above this size, the predators will only injure them or even fail to catch them. Larvae laid in autumn reached this limit size in about 3 months, whereas only 1.5 month is necessary for larvae laid in spring (personal observation). Thus, we used the mean density of predators and competitors over a period of 3 months after spawning date for autumn tadpoles and 1.5 months for spring tadpoles.

### Statistical analyses

All statistical analyses were performed on R 3.4.1 (R Core Team, 2018). To assess if pond characteristics differed between seasons, we applied a linear model for the depth of the pond and generalized linear models with a quasi-Poisson family for all other variables to account for overdispersion. The *presence of egg masses* and the *number of egg masses* were analysed using a generalized mixed model with site as random effect, with a binomial family or a negative binomial family (to account for overdispersion), respectively. The hatching rate and survival rates from egg to metamorph were often zero hence we decided to analyse them as binary variables using a generalized mixed model with site as random effect and a binomial family. Those derived binary variables, called *hatching success* and the *metamorphosis success* are the probability of observing at least one hatchling or one metamorph in a pond where egg masses were observed. The significance of fixed effects were tested using Chi^2^ tests to compare nested models (Zuur et al., 2009).

### Bet-hedging model

Finally, we wondered if the coexistence of two breeding periods could result from a bet-hedging strategy, with the optimal strategy being to split the breeding effort between the two favourable seasons to spread the risk of complete failure (Seger & Brockman, 1987). The following model is an ESS model derived from Saiah and Perrin (Saiah & Perrin, 1990) on the hatching probability of fairy shrimp seasonal cohorts. Their model was primarily inspired by Cohen (1966), on the optimal reproduction strategy of an annual plant whose seeds can either germinate or remain dormant. In our case, there are two strategies: autumn breeding with initial success (i.e. the ability of offspring to persist until spring) depending on the environmental conditions, and spring breeding with success depending mainly on the presence of autumn tadpoles, hence on the initial success of autumn breeding (as suggested by the results on success of autumn and spring breeding events, see below).

Let *c* be the proportion of eggs laid in autumn (thus 1-*c* in spring) – we assume, in agreement with our data (see results), that *c* represents a fixed strategy, i.e. the frogs cannot predict failure in advance to avoid laying in autumn, nor can they avoid laying eggs in spring when an autumn cohort is present. As mentioned above, the autumn cohort is assumed to succeed or fail, at random, with probability *q* and 1-*q* respectively. When it succeeds, a fraction *s_1_* of the offspring survive to reproductive age. The spring cohort completely fails whenever the autumn cohort has survived in a pond (a reasonable simplification based on our survival rates estimates, see below), otherwise a proportion *s_2_* of spring tadpoles survive. Overall, the expected number of offspring reaching sexual maturity is *c* s_1_ when the autumn cohort doesn’t fail and (1 – *c*) s_2_ when it does.

If each frog reproduced only during one year, the optimal strategy would maximize the geometric mean of the annual reproductive outcome (Dempster, 1955) which is

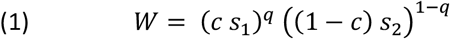

Or, equivalently

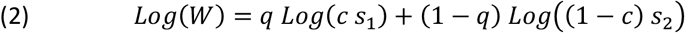

However, reproductive life lasts more than one year in frogs (say, *n* years), which in itself is a way to spread the risk of failure among successive cohorts of offspring – an uncertainty remains however, for each frog, on how many (*k*) of the *n* breeding years will not allow the autumn cohort to survive. For each individual, *k* is distributed binomially with probability 1-*q* so that

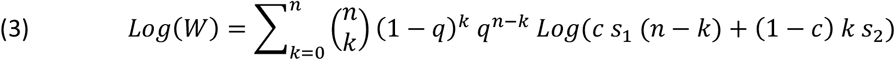

where 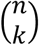 represents the number of possible repartitions of the k years with autumn failure among the total number of breeding years *n*.

The selection gradient on *c* is the derivative of the function Log(W), which indicates whether selection favours an increase in *c* (if positive) or a decrease (if negative):

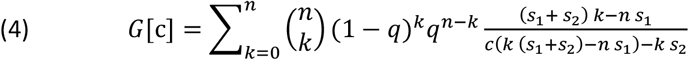

If some value of *c* within the authorized interval [0,1] results in *G*[c]=0 then it is considered an evolutionary stable strategy (ESS) provided the second derivative is negative (i. e. *G*[c] is positive below the ESS and negative above).

We explored numerically the selection gradients in order to find potential ESS using Mathematica (Wolfram Research Inc., 2018) based on the following parameter combinations. We set survival probabilities based on our estimates of survival from egg to metamorphose: *s_1_* = 0.047 (estimated among breeding events producing offspring that survived until spring) and *s_2_* = 0.038 (in the absence of autumn tadpoles). We assumed that survival and fecundity were equal for both seasonal cohorts for the rest of the life cycle. We set the number of reproductive years *n* = 3 to 5, according to a study of age structure of a breeding population in Spain (Esteban et al., 2004). Note that this model applies at the individual level (as developed above) as well as at the genotype level.

## Results

### Characteristics of temporal niches

Pond depth was not significantly different between the autumn (here from September to December) and spring (here from January to April) breeding seasons (Table 1). The densities of amphibian larvae (other than Parsley Frog) were not significantly different. In autumn, extreme densities of *Epidalea calamita* tadpoles were recorded in some small ponds whereas the well-known spring breeding-species (*Hyla meridionalis, Pelophylax sp., Triturus marmoratus, Lissotriton helveticus*) reproduce later than the Parsley Frog, hence their larvae are only present from April onwards. The density of potential invertebrate predators was higher in autumn than in spring (χ^2^_1_ = 37.17, p-value = 0.005) with the lowest density being from December to March. On the contrary, the number of adult newts (potential predators of Parsley Frog tadpoles) was higher in spring than in autumn (χ^2^_1_ = 369.36, p-value = 2.2*10^−16^).

**Table 1:**
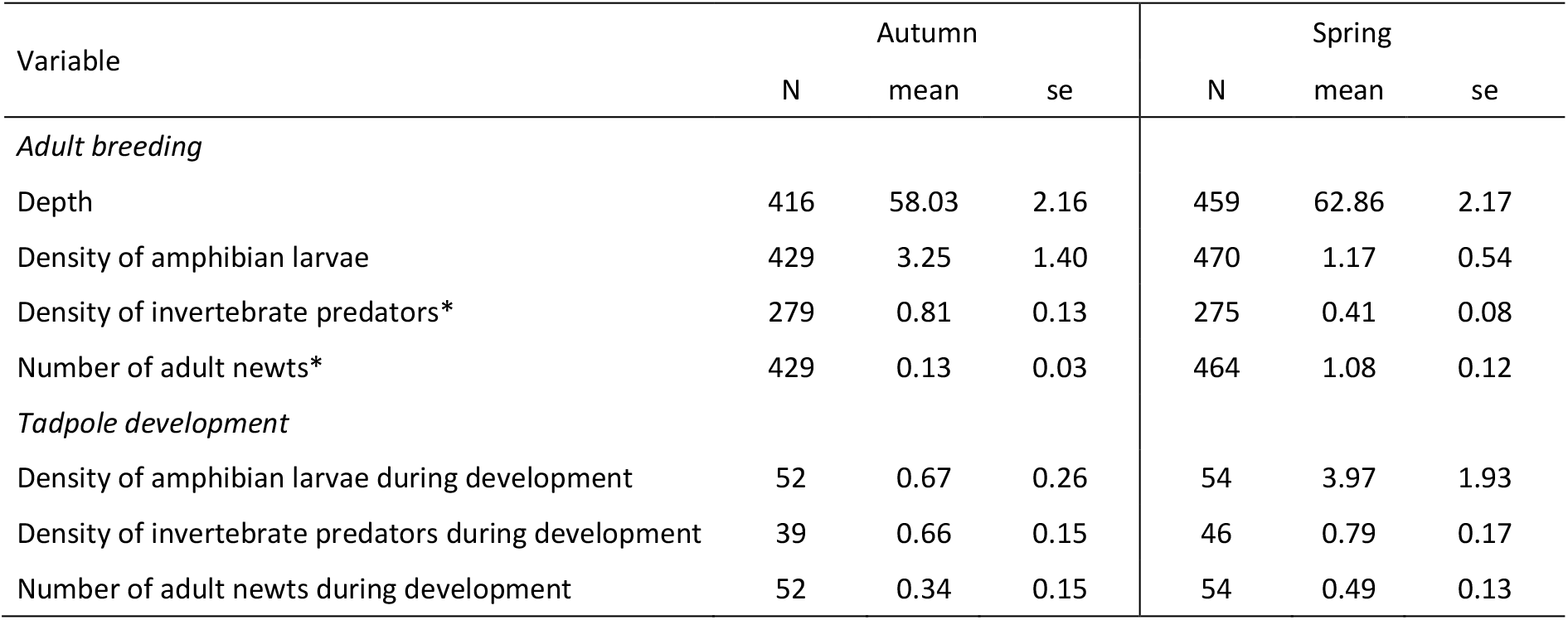
Description of the seasonal niches regarding adult breeding (upper part) and larval development (lower part) of the Parsley Frog. Mean and standard error (se) of the explanatory variables depending on the season (autumn and spring). N: sample size. *P.p* is *Pelodytes punctatus*. Depth is in centimeters. Density of amphibian larvae or invertebrate predators is the mean number of individuals sampled in one dipnet sweep. Parsley Frog is excluded from calculations indicating “amphibian larvae” or “anuran adults”. * denotes significant difference between season for the considered variable.

### Breeding phenology

We registered 184 breeding events, 79 in autumn and 105 in spring. Note that in two sites, one breeding event was recorded in May. The *number of egg masses* recorded in one pond showed a bimodal pattern with a peak in October and another in February (Fig. 1).

**Figure 1:**
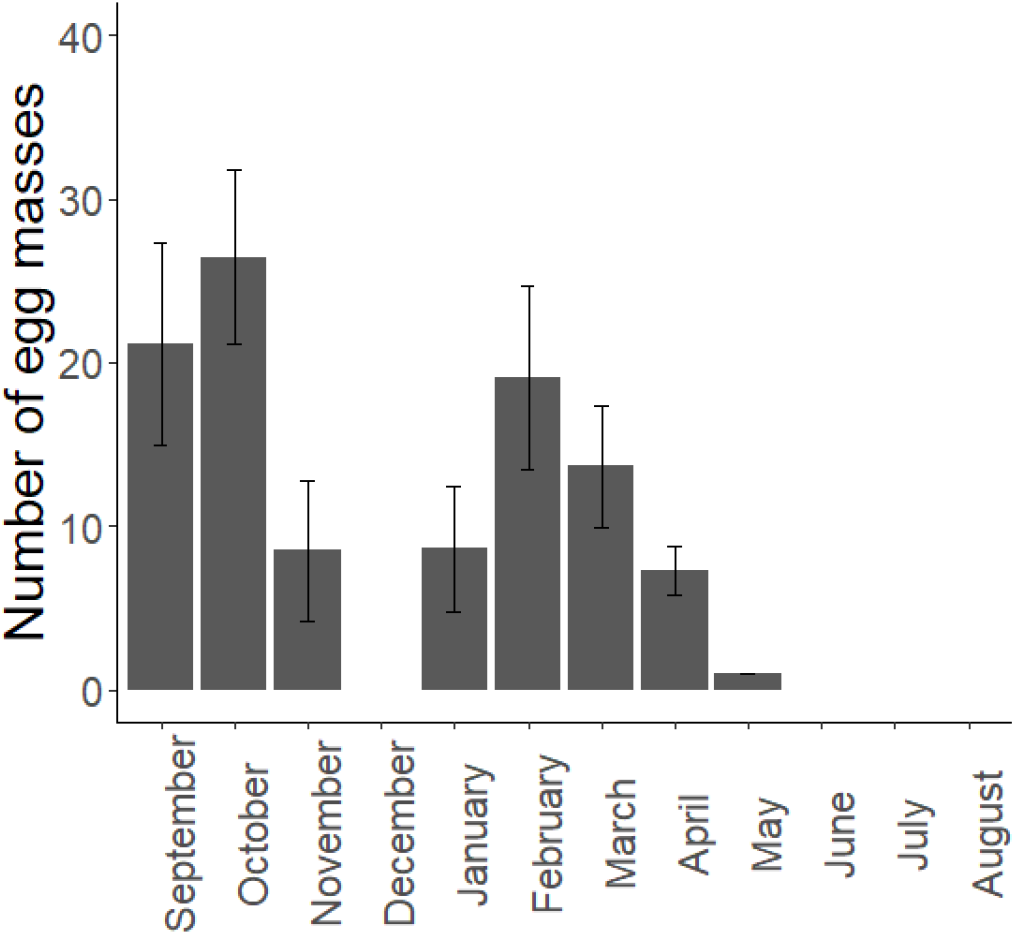
Mean monthly number of egg masses produced by the Parsley Frog for each recorded breeding event. Error bars are standard errors among sites among years.

The *presence of egg masses* (finding at least one egg mass when visiting a pond) was not significantly different between the two seasons (0.18 ± 0.02, mean ± S.E per visit in autumn and 0.22 ± 0.02 per visit in spring, (over 429 and 470 visits, respectively) χ^2^_1_ = 2.31, p-value= 0.128, see also Table 2). This variable was not affected by the presence of anurans from other species (larvae), nor by the presence of predators (invertebrates or adult newts). It was positively related to the depth of the pond (χ^2^_1_ = 20.40, p-value = 6.3*10^−6^). The *presence of egg masses* observed in spring was not affected by the presence of autumn tadpoles (χ^2^_1_ = 0.03, p-value = 0.875).

**Table 2:** Results of the statistical analyses performed to explain the variation of *presence of egg masses, number of egg masses, hatching success* and *metamorphosis success*. Bold letters indicate a significant test (p-value < 0.05).

| Variable | Tested effect | $\chi^2$ value | p-value |
| --- | --- | --- | --- |
| Presence of egg masses | Season | 2.3101 | 0.1284 |
|  | Depth | **20.398** | **6.289e-06** |
|  | Density of invertebrates | 0.1869 | 0.6655 |
|  | Number of adult newts | 0.2419 | 0.6229 |
|  | Density of amphibian larvae | 0.1292 | 0.7192 |
|  | Number of adult anurans | 0.3972 | 0.5286 |
| Number of egg masses | Season | **9.25** | **0.0023** |
| Hatching success | Season | **11.119** | **0.00085** |
|  | Density of invertebrates | 0.1549 | 0.6939 |
|  | Number of adult newts | 0.8315 | 0.3618 |
|  | Density of amphibian larvae | 0.1501 | 0.6984 |
| Metamorphosis success | Season | 0.3932 | 0.5306 |
|  | Density of invertebrates | 0.6914 | 0.4057 |
|  | Number of adult newts | 0.4911 | 0.4835 |
|  | Density of amphibian larvae | 1.8085 | 0.1787 |

The *number of egg masses* was higher in autumn than in spring (23.0 ± 4.0 egg masses per breeding event in autumn and 13.7 ± 2.4 in spring; χ^2^_1_=9.25, p-value = 0.002, Table 2, Fig. 2.A and Annex 2). As a result, autumn breeding contributed slightly more than spring breeding to the production of egg masses (57% versus 42.9%).

**Figure 2:**
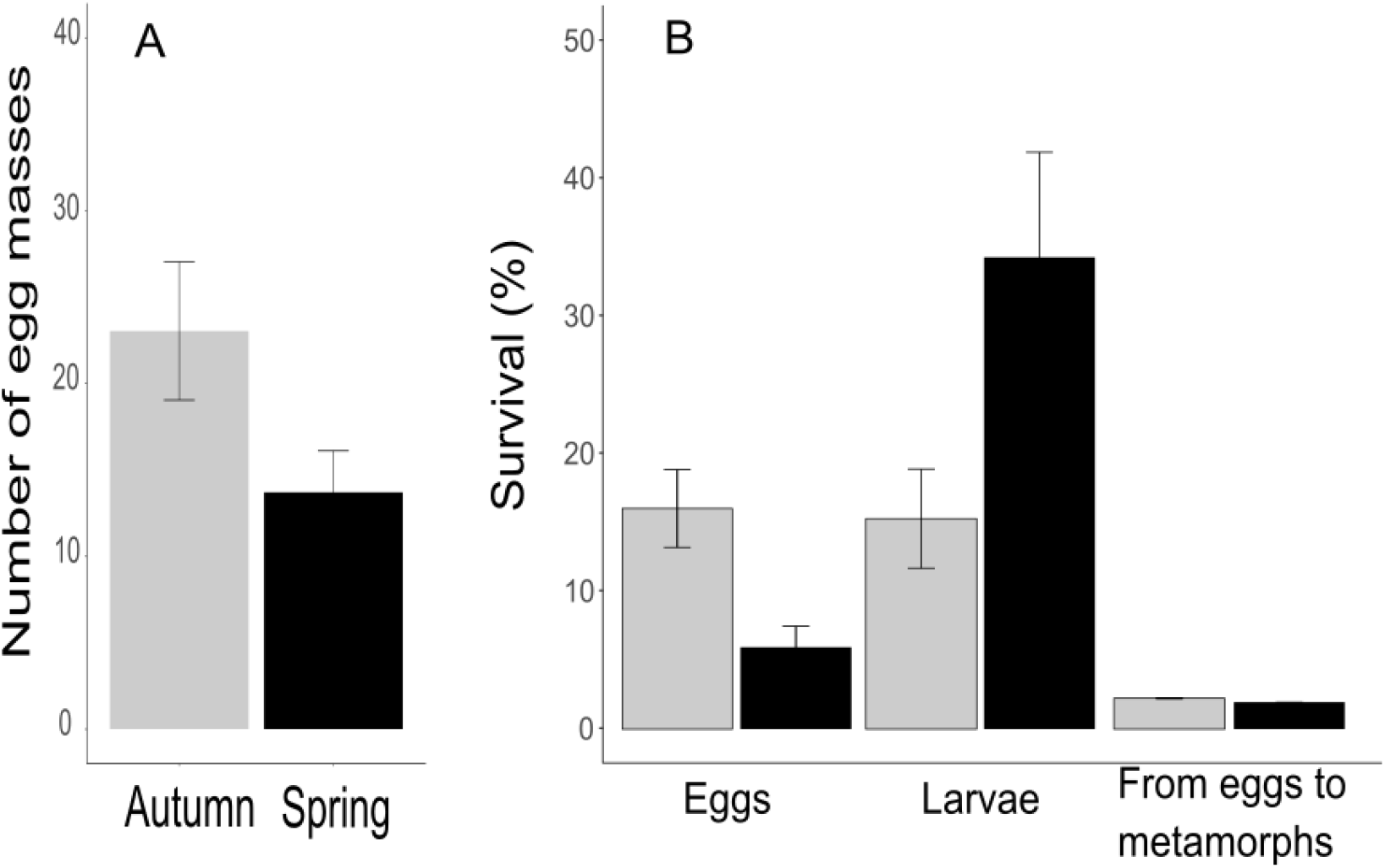
**A** Mean number of egg masses produced per season by the Parsley Frog for each recorded breeding event. **B** Mean survival rates during embryonic stage (hatching rate, n = 159), larval stage (n = 79) and from eggs to metamorphs (n = 163). Error bars are standard errors, among sites, among year. Autumn in grey and spring in black.

### Breeding success

*Hatching success* (i.e. the percentage of breeding events producing at least one larvae) was higher in autumn than in spring (68.4% and 43.8% respectively, χ^2^_1_ = 11.12, p-value = 0.001, Table 2). *Metamorphosis success* (i.e. the percentage of breeding events producing at least one metamorph) was not significantly different between the two seasons (34.2% in autumn and 29.8% in spring χ^2^_1_ = 0.39, p = 0.531).

Drought (pond totally dried up) caused the total failure of 7 breeding events in autumn and of 5 breeding events in spring over the 3 year-survey and the 19 sites (representing 9% and 4.8% of the breeding events, those percentages are not significantly different, χ^2^_1_ = 0.66, p = 0.42). Drought caused mortality of offspring at different developmental stages (mostly eggs for autumn cohort and tadpoles for spring cohort). Neither *hatching success* nor *metamorphosis success* were explained by interspecific competition (the density of other amphibian larvae) or by predation (density of potential invertebrate predators or number of adult newts, see Table 2).

Survival rates are represented in Figure 2.B and Annex 3. The *survival rates from egg to metamorph* were similarly low (autumn: 2.24 % ± 0.61 and spring: 1.97 % ± 0.73, Table 2), resulting in a higher contribution (74.6%) of autumn breeding in the overall production of metamorphs per site and per year (due to the higher breeding effort in autumn, see above).

The autumn cohort persisted until spring in 34/79 breeding events (43%, corresponding to the rate of initial success, *q*, see bet-hedging model). In those cases, tadpoles laid in spring coexisted in their pond with tadpoles from the autumn cohort. The presence of an autumn cohort of Parsley Frog tadpoles significantly reduced the *metamorphosis success* of spring cohorts to 18.4% in presence of autumn tadpoles, versus 40.0% in absence of autumn tadpoles, χ^2^_1_ = 10.60, p-value = 0.005).

This reduction effect was not significant for the *hatching success* (32.6% in presence of autumn tadpoles, versus 53.6% in absence of autumn tadpoles χ^2^_1_ = 4.63, p-value = 0.099). Accordingly, all three survival rates were reduced in the presence of autumn tadpoles and this effect was most pronounced for the *survival from egg to metamorphs* (3.77% ± 1.4 versus 0.16% ± 0.08 in absence versus presence of autumn tadpoles, Fig. 3 and Annex 4). We also tested if the *metamorphosis success* of autumn tadpoles might be affected by the presence of spring tadpoles, but this was not the case (χ^2^_1_ = 2.75, p-value = 0.097).

**Figure 3:**
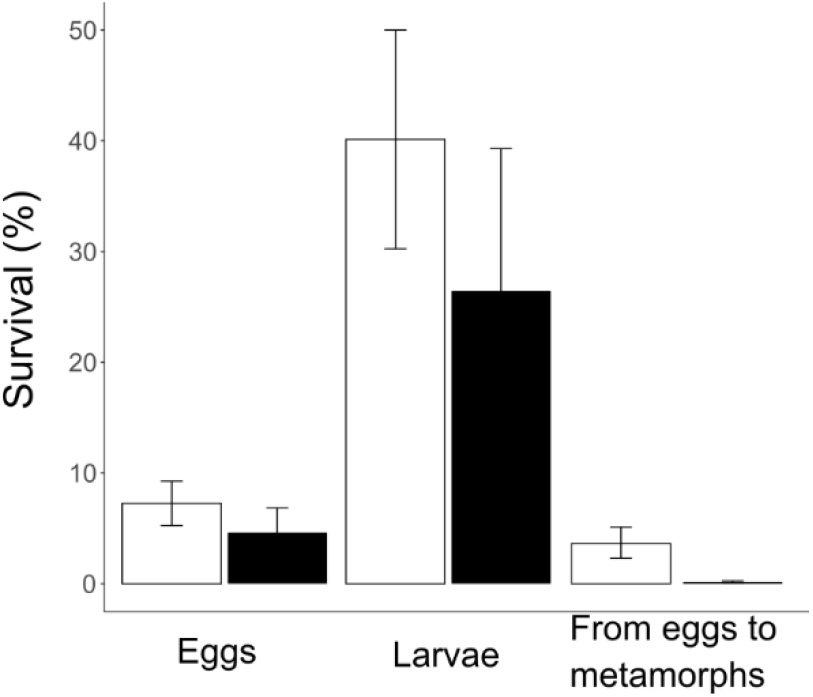
Mean survival rates during embryonic stage (hatching rate n = 86), larval stage (n = 27) and from eggs to juveniles (n = 90) of spring cohorts, in presence (black) or absence (white) of older tadpoles laid in autumn. Error bars are standard errors, among sites, among year.

Finally, Figure 4 summarizes the breeding strategy of Parsley Frog showing the presence of egg masses, tadpoles and the metamorphs in each studied site, over the three years of survey. It illustrates the quasi-exclusion between the two cohorts: there were only 4 cases in total (out of 47 observations) where metamorphs from the two seasonal cohorts emerged in spring in the same pond during the same year.

**Figure 4:**
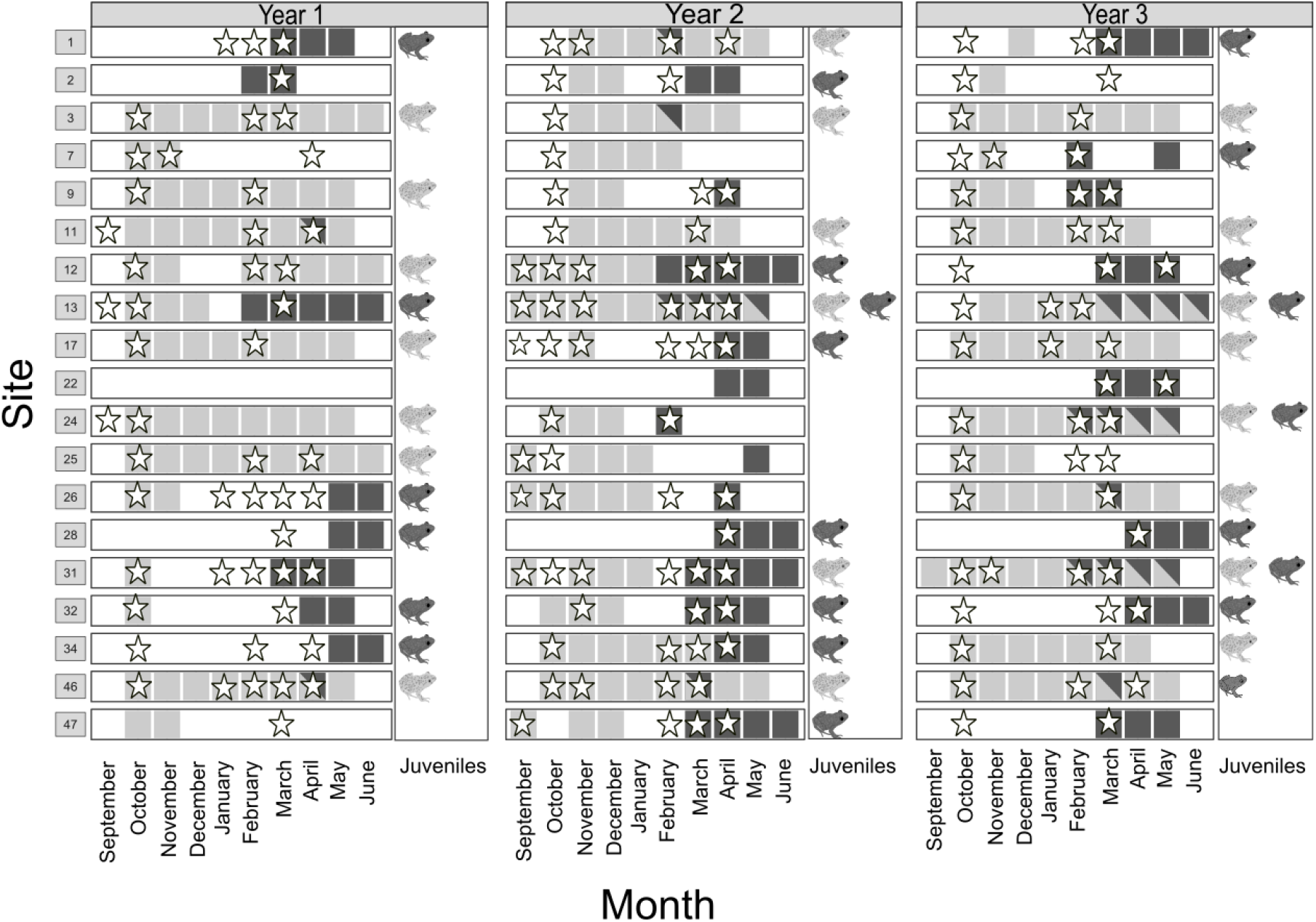
Summary of breeding strategy of the Parsley Frog in the 19 studied sites (vertical lines) in the three successive years: presence of egg masses (stars), presence of tadpoles (squares) and presence of metamorphs (frogs). Grey is indicative of autumn events and black is indicative of spring events.

### Maintenance of spring breeding

Selection gradients based on our bet-hedging model predict that a mixed strategy is maintained when the rate of initial success of the autumn cohort (*q*) is between 0.2 and 0.8 for a number of reproductive years n = 3. In this condition, a pure autumn strategy is predicted above 0.8, and a pure spring strategy below 0.2. (Fig. 5). The maintenance of this strategy is less probable if the number of reproductive years increases (n = 5 years of breeding), with a reduced range of *q* leading to a stable mixed strategy.

**Figure 5:**
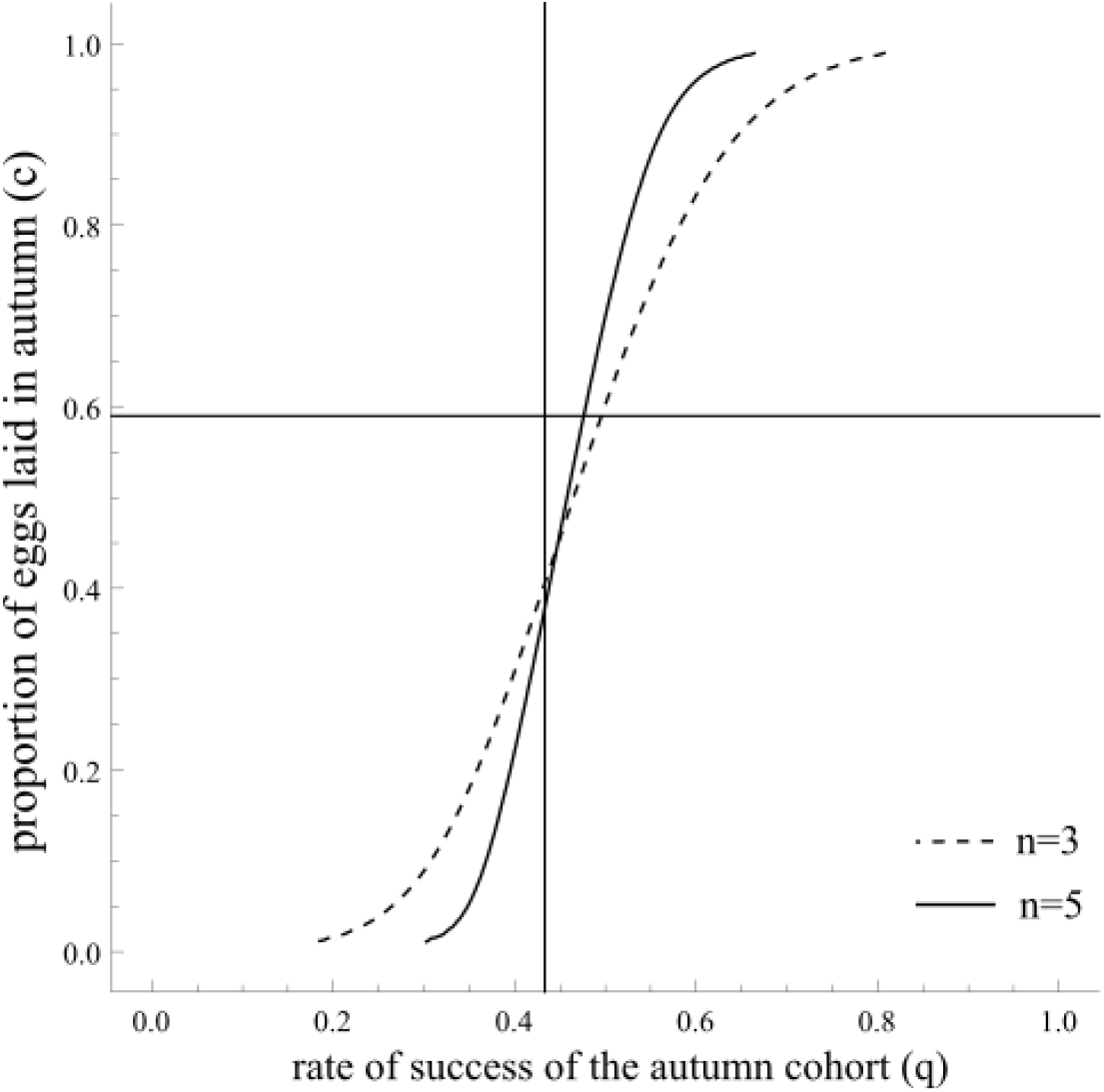
Evolutionary stable strategy (black lines), based on a bet-hedging model, predicting the proportion of eggs laid in autumn (*c*, x-axis) in relation to the rate of success of the autumn cohort (*q*, y-axis), depending on the number of breeding years (n). We set the survival rate of autumn tadpoles to 4.7% (estimated among breeding events producing offspring that survived until spring) and the survival rate of spring tapdoles to 3.8%, (estimated in absence of autumn cohort). The horizontal and vertical lines indicate the field estimates of *c* and *q*.

## Discussion

### Cost and benefits of a bimodal breeding phenology

We used field surveys to describe the breeding phenology of the Parsley Frog in the French Mediterranean region but also to quantify the relative contribution and success of each seasonal reproduction (autumn and spring reproduction). This quantification, rarely achieved in the wild (but see Licht 1974; Banks & Beebee 1988; Gascon 1992; Wheeler *et al*. 2015), is essential to understand the evolution of this bimodal breeding strategy. We confirmed the existence of two distinct seasonal peaks in breeding activity, probably mediated by cold temperature in December and January as adult Parsley Frogs tend to breed in mild and rainy periods as was previously observed (Toxopeus et al., 1993; Guyétant et al., 1999; Jakob et al., 2003). However breeding episodes occurred even in the absence of rainfall as long as ponds were filled with water (personal observations and Richter-Boix *et al*. 2006b).

The *number of egg masses* in our population was higher in autumn than in spring. This is in apparent contradiction with Richter-Boix et al. (2006b) who found that a four-fold higher breeding effort in spring than in autumn in the northeast of the Iberian Peninsula. While we don’t have a definitive explanation for this difference, we suggest it could be related to higher competition among anuran larvae in autumn in north-east Spain compared to France. In our study area in southern France, larvae of *Pelodytes punctatus* are typically the only anuran larvae found after the summer drought in the ponds in autumn. In contrast, four other species of Anura (*Hyla meridionalis, Epidalea calamita, Alytes obstetricans* and *Pelophylax perezi*) have tadpoles in autumn and three of them (i.e. all except *E. calamita*) can have overwintering tadpoles in Spanish ponds (Richter-Boix et al., 2006a). These authors also showed that *Pelodytes punctatus* tadpoles suffer from interaction with *Hyla meridionalis* (Richter-Boix et al., 2007b). It is thus possible that increased competition for *Pelodytes punctatus* larvae in autumn and winter makes the autumn niche less favourable in north-eastern Spain compared to our study area in southern France and reduce Parsley Frog investment in autumn breeding there.

The *survival rate from egg to metamorph* was low in both seasons. The combination of *numbers of egg masses* and survival rates eventually resulted in a higher contribution of autumn breeding to the overall production of metamorphs. The overall low survival rates of offspring that we found is in line with previous field studies in anurans (e.g. Licht 1974; Banks & Beebee 1988) and can be caused by pond desiccation, predation, inter and intra-specific competition for food and parasitism or pathogen infections. Our study revealed no obvious effect of variation in predation on tadpole survival even if the predation pressure encountered by tadpoles at the beginning of their development varies from site to site (but not between seasons). This may seem surprising since many studies experimentally demonstrated that predation cause substantial mortality to tadpole populations (e.g. Tejedo 1993; Van Buskirk & Arioli 2005). This may be due to the lack of information about predation during the first year of survey which reduced our statistical power or to the fact that causative factors are numerous and more complex to identify in the field. However, other studies reported no effect of predation on tadpole survival (Hartel et al., 2007) or even a positive effect (Barandun & Reyer, 1997), probably due to predator-induced phenotypic plasticity. Nevertheless, our results suggest that the predation pressure is probably not a stronger constraint in one season than in the other.

Spring tadpoles should be exposed to more competitors during their development than autumn tadpoles since the majority of amphibian species in the local community breed in March and April. Nevertheless, we found no effect of interspecific competition on survival for any of the two seasonal tadpole cohorts. This seems surprising since Parsley Frog is a poor competitor as a tadpole compared to most species of the anuran community, in particular *Hyla meridionalis* and *Rana perezi*, present in spring in permanent ponds (Richter-Boix et al., 2007b). On the contrary, in small temporary ponds and during autumn and winter season, Parsley Frog tadpoles encounter mostly *Bufo bufo* and *Epidalea calamita* with even lower competitive abilities (Richter-Boix et al., 2007b). We hypothesize that interspecific competition effect was not detected in our study due to numerous uncontrolled sources of variation.

### Priority effects

We revealed a striking negative effect of the presence of conspecific autumn tadpoles on the survival of spring tadpoles in the Parsley Frog. Previous studies have demonstrated the occurrence of such intraspecific priority effects in amphibians in experimental settings (Morin et al., 1990; Eitam et al., 2005; Murillo-Rincón et al., 2017) but as far as we know, our study is the first evidence for intraspecific, inter-cohort competition in amphibians in nature. In the field, we observed in several occasions that large autumn tadpoles were eating freshly laid eggs of their own species, which could partly explain the lower hatching rate of spring eggs in presence of autumn tadpoles. Moreover, (Tejedo, 1991) previously described how Parsley Frog tadpoles preys on *Epidalea calamita* eggs. In this latter study, predatory tadpoles were exclusively old tadpoles and they could cause a loss of 50 to 100% of the eggs. Oophagy has also been demonstrated to be responsible for interspecific priority effects between *Scaphiosus couchii* and *Bufo speciosus* (Dayton & Fitzgerald, 2005). Intraspecific oophagy has been described in some anuran species (Summers, 1999; Dayton & Wapo, 2002) and has been proposed as an energetic opportunistic response in food shortage in temporary ponds.

However, the presence of autumn tadpoles also affects the larval survival (post-hatching) of spring tadpoles. This may reflect competition for resources between large autumn and small spring tadpoles as shown in *Rana arvalis* (Murillo-Rincón et al., 2017). Interference competition mediated by microorganisms may also play a role: smaller tadpoles could display coprophagy instead of feeding on higher quality resources (Beebee & Wong, 1992; Baker & Beebee, 2000). This large priority effect between the two seasonal tadpole cohorts of Parsley Frog has a great impact on the overall efficiency of breeding: in most ponds, there could be only one successful breeding period, autumn or spring. Nonetheless, we found no indication that spring breeders select their oviposition site to avoid conspecifics, as other amphibian species sometimes do (Sadeh et al., 2009). Accordingly, the *presence of egg masses* was also unaffected by the presence of potential competitors or predators.

### Seasonal partitioning of breeding: a bet-hedging strategy?

The temporal partitioning of breeding activity could reflect several evolutionary processes: 1) the existence of two specialized phenotypes either genetically determined (in which case we would expect temporal genetic differentiation between cohorts) or set by early environmental cues (phenotypic plasticity); 2) a use of alternative strategies by some or all individuals (bet-hedging). We previously demonstrated that the two temporal cohorts do not reflect two genetically distinct temporal populations (Jourdan-Pineau et al., 2012) but breeding phenology may still be set once for good for each individual. In this case, breeding in autumn or in spring could be determined by the physiological state (and sexual maturity) of the breeder and maintained year after year, by physiological constraints (typically the case for a capital breeder species which stores energy for future reproduction (*e.g*. Lardner & Loman 2003). In a diversified bet-hedging strategy, individual breeding activities could vary from year to year (each year, individuals would “choose” one breeding season) or individuals could split their breeding effort between the two seasons in some or most years. There is no individual data available for this species and our only attempt to mark adults with visible implant alpha tags was not successful. Preliminary results based on genotyping of eggs, spawned in the same pond at different periods, suggests that females could breed several times in one year but this has to be confirmed (unpublished data). Clearly, this is a line of research to develop in the future if we want to fully understand the evolution of reproduction in this system.

Based on our field survey, it appears that the bimodal breeding phenology of Parsley Frog is a typical diversified bet-hedging strategy. The large priority effect between the two seasonal cohorts, combined with high unpredictability of conditions that result in failure or success of entire cohorts, results in frequency dependent-selection and favour risk-spreading strategies: the best option is to develop in ponds with the smaller number of conspecific competitors. These conditions are found partly in autumn, when the habitat becomes favourable after the dry summer period, or in spring, as some of the autumn cohorts have died in the winter, leaving the habitat free. Poethke et al. (2016) developed a theoretical model in which they outlined this impact of competition on the evolution of bet-hedging strategy. Using a model for optimal germination fraction, based on field data on desert plants, Gremer and Venable (Gremer & Venable, 2014) also showed that density-dependence could explain the observed bet-hedging strategy of germination spread in time (i.e. not all seeds at once). Density-dependence was not included in our model and we do not have field data to assess its effect in our populations. This would be a fruitful line of research to improve our understanding of this breeding system.

In the Parsley Frog, our model shows that the observed mixed breeding strategy (0 < *c* < 1) is maintained if the rate of initial success of the autumn cohort (*q*) is between 0.2 and 0.8 (if females have on average 3 years of breeding in their lifetime) or between 0.35 and 0.65 (for 5 years of breeding). Those conditions are fulfilled according to our field estimates (*q* = 0.43). We estimated the proportion of eggs laid in autumn by all breeders (*c* = 0.57) but could not estimate this proportion at the individual level. Survival rates set in the model were based on our field estimates of survival from egg to metamorphosis; hence, we assumed similar survival after metamorphosis of the two cohorts. Unfortunately, we have no information about survival of Parsley Frog during its adult terrestrial life. However, the adult survival is an important parameter in our model since it determines the number of reproductive years. The mixed breeding strategy is less stable when the number of breeding opportunities per lifetime increases – as the risk is now spread over several successive years. Indeed, experiencing variation in reproductive success among those opportunities is less harmful when it is possible to try again the next year. A skeletochronology study conducted in a upland population in Spain indicated that the mean age of sampled Parsley Frog females was 5.01 years (with a standard deviation of 1.99) (Esteban et al., 2004). Assuming a minimal age at first reproduction of 1 year (as done by Esteban *et al*. 2004), this translates into an average number of reproductive years or females of n = 4. Our evaluation of the bet-hedging strategy with n = 5 is thus probably conservative.

We previously showed that the Parsley Frog successfully exploits two temporal niches in the Mediterranean region thanks to a high phenotypic plasticity of tadpole development to face very different seasonal environments (Jourdan-Pineau et al., 2012). Recently, the combination of phenotypic plasticity and bet-hedging has been theoretically investigated, suggesting that phenotypic plasticity could further minimize fitness variances caused by mismatches between phenotype and environment (Rádai, 2020; Haaland et al., 2021). Interestingly, in the wolf spider, temperature and day length leads to alternative developmental types within broods. This cohort splitting is both probabilistic and sensitive to environment, a phenomenon proposed as being a plastic bet-hedging strategy by Rádai (2020). In this case, the various plastic phenotypes, triggered by environmental variations, constitute a bet-hedging response to grassland habitats with substantial and unpredictable year-to-year variation.

In the Parsley Frog, priority effects are not the only factor influencing the relative success of the spring and autumn strategies in terms of future recruitment: autumn tadpoles metamorphose earlier and at a much larger size than spring tadpoles (Jourdan-Pineau *et al*. 2012, unpublished data). This should confer to them a significant advantage in terms of survival to adulthood (Smith, 1987; Altwegg & Reyer, 2003; Székely et al., 2020) even if we don’t know whether size and date of metamorphosis affects survival and ultimately fitness in our model. In addition, there is no significant difference in cohort survival (the probability to produce at least one metamorph) between spring and autumn, in spite of a slightly higher risk of drought (and hence complete disappearance of the cohort) in autumn. Density-dependence (on which we have no information) might partly explain why autumn cohorts do as well as spring cohorts in spite of higher drought risk. Additionally, our measures of breeding success are very rough because counting precisely the number of larvae from each cohort in the ponds over the course of the season is extremely difficult. There is thus still much to learn to fully understand the advantages and disadvantages of autumn and spring strategies in this species.

Lastly, as explained above, we still don’t know if individual females usually breed once a year (either spring or autumn) or several times a year (potentially spring and autumn of the same year). Capture-mark-recapture of adults and larvae would alleviate some of these limitations but would be highly challenging. However, our results remain valid for a large range of parameters, and these uncertainties should not affect our conclusion that the breeding strategy of Parsley Frogs in southern France constitutes an original example of a bet-hedging strategy driven by high environmental stochasticity and large inter-cohort priority effect.

## Acknowledgements

We are grateful to Vincent Mouret, Alain Fizesan, Denis Rey, Simon Russeil and Jérémy Aubain for help in field work. Virginie Ravigné provided valuable help with the bet-hedging model.

Preprint version 5 of this article has been peer-reviewed and recommended by Peer Community In Evolutionary Biology (https://doi.org/10.24072/pci.evolbiol.100147).

## Data, scripts and codes availability

Data and scripts are available online: https://doi.org/10.18167/DVN1/2TOXXV.

## Appendix

**Annex 1:** Surveyed ponds (n°, names and area) and their geographic localization (in decimal degrees, WGS84).

| Site # | Pond name | Area | Latitude | Longitude |
| --- | --- | --- | --- | --- |
| S01 | Pataris | Cournonterral | 43.5694 | 3.6951 |
| S02 | Fertalière | Cournonterral | 43.569 | 3.6812 |
| S03 | Logis du Bois | Notre-Dame de Londres | 43.8406 | 3.7462 |
| S07 | Jasses | Notre-Dame de Londres | 43.8657 | 3.7828 |
| S09 | SEGA | Notre-Dame de Londres | 43.8147 | 3.7817 |
| S11 | Fesq | Notre-Dame de Londres | 43.8028 | 3.7919 |
| S12 | Bouillés Ouest | Cournonterral | 43.5694 | 3.6951 |
| S13 | Bergerie | Cournonterral | 43.5803 | 3.7019 |
| S17 | Combe l'Escure | Cournonterral | 43.5733 | 3.6994 |
| S22 | Pioch noir | Gardiole | 43.4947 | 3.7544 |
| S24 | GMP | Cazevieille | 43.7708 | 3.7945 |
| S25 | PMP | Cazevieille | 43.7708 | 3.7950 |
| S26 | Seuilles | Cazevieille | 43.7597 | 3.7772 |
| S28 | Anduze | Gardiole | 43.4787 | 3.7318 |
| S31 | Farlet | Gardiole | 43.4908 | 3.7415 |
| S32 | Pierre Tintante | Gardiole | 43.4826 | 3.7341 |
| S34 | Roubiac | Cazevieille | 43.7597 | 3.7879 |
| S46 | Boissière | Notre-Dame de Londres | 43.8337 | 3.7461 |
| S47 | Mège | Gardiole | 43.4936 | 3.7544 |

**Annex 2:**
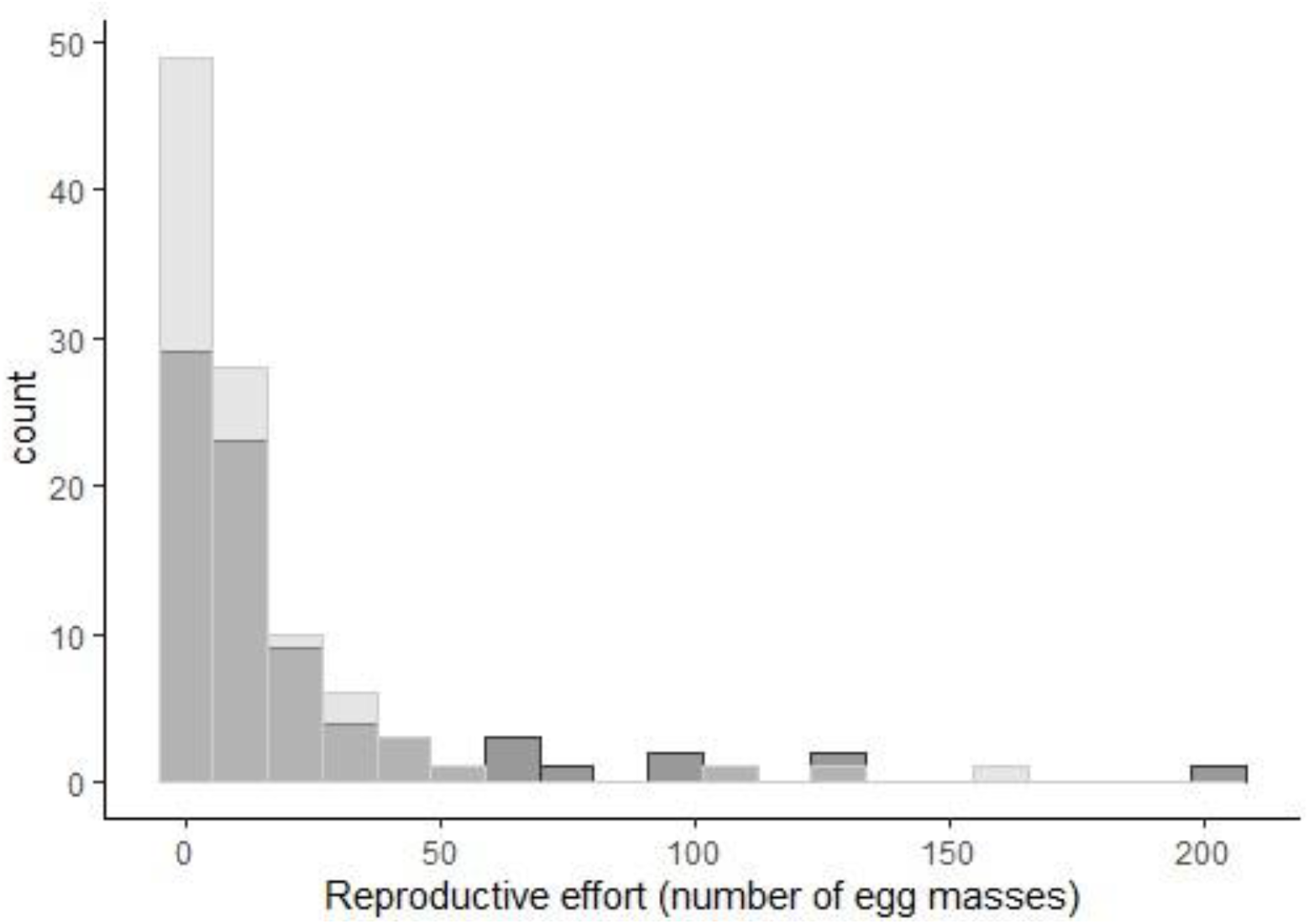
Distribution of the *number of egg masses* produced at each breeding event per season. Autumn in dark grey and spring in light grey.

**Annex 3:**
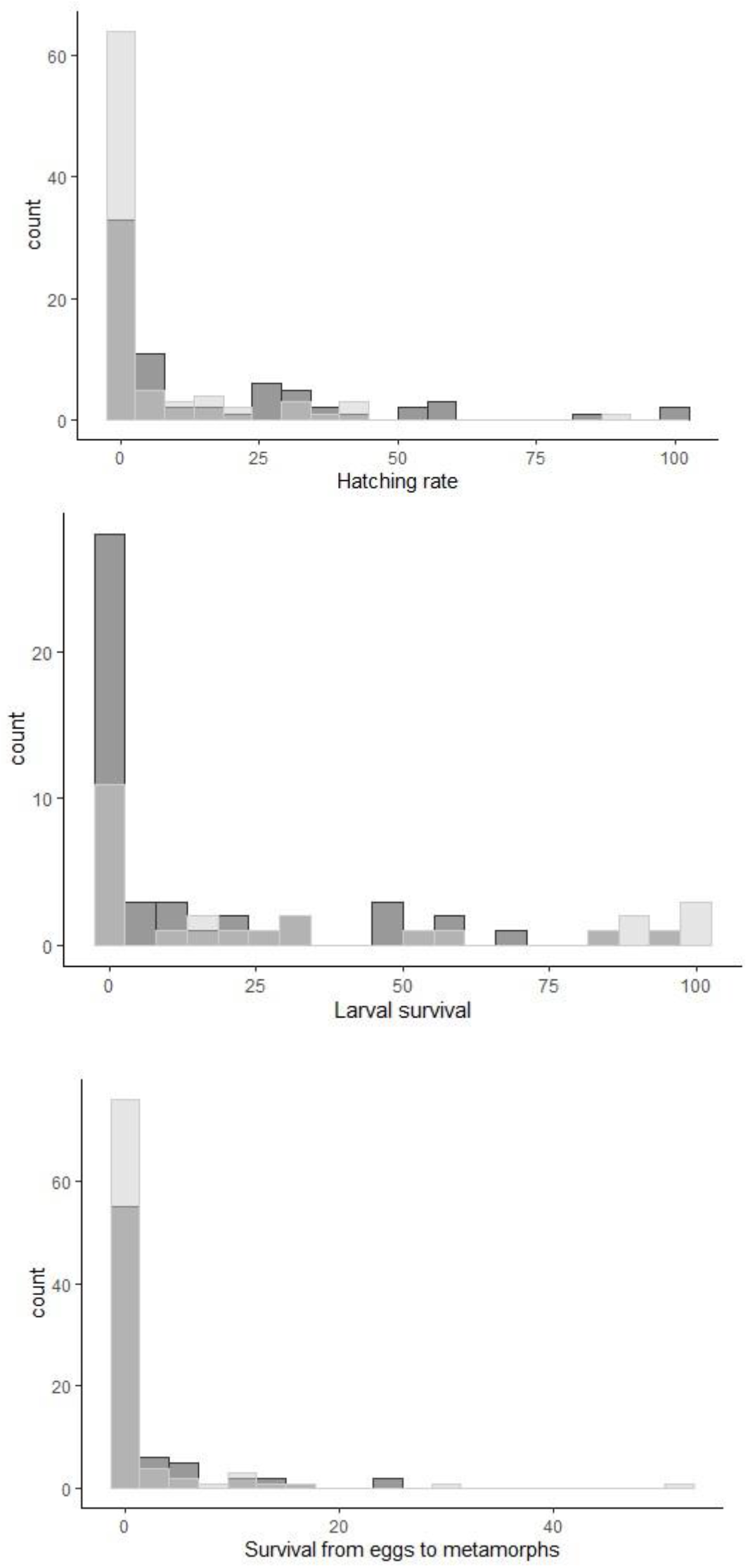
Distribution of hatching rate, survival rate during larval stage and survival rate from eggs to metamorphs. Autumn in dark grey and spring in light grey.

**Annex 4:**
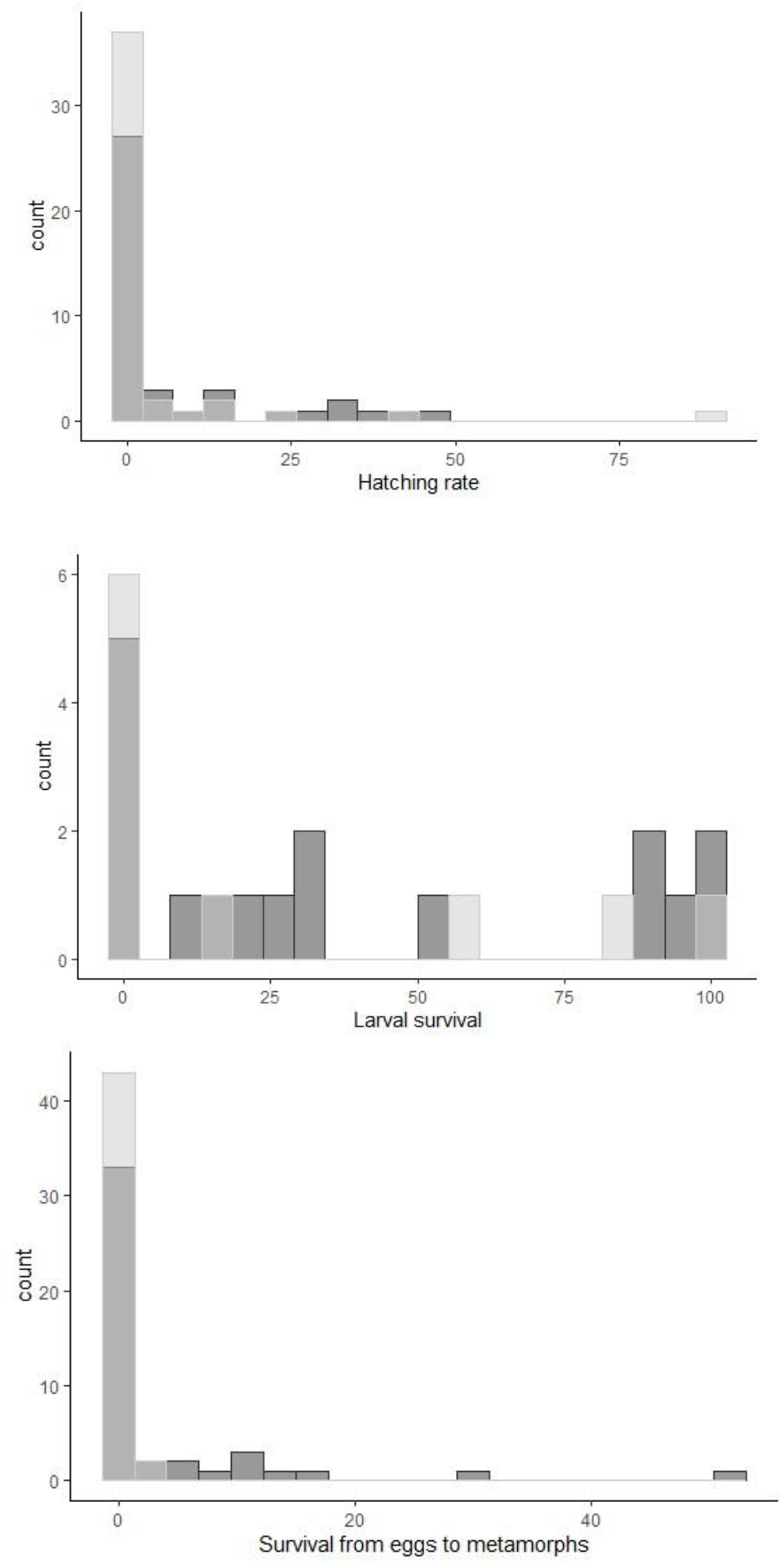
Distribution of *hatching rate, survival rate during larval stage* and *survival rate from eggs to metamorphs* of spring cohorts, in presence (dark grey) or absence (light grey) of older tadpoles laid in autumn.

